# Negatively charged α-Synuclein condensate modulates partitioning of molecules

**DOI:** 10.1101/2024.07.24.604760

**Authors:** Qingqing Yang, Shunfa Chen, Pengfei Zhang, Zhonghua Lu, Shuwen Chang, Leo E. Wong

## Abstract

α-Synuclein (αSyn) aggregation via liquid-liquid phase separation has recently emerged as a crucial mechanism underlying amyloid fibril formation implicated in Parkinson’s disease. However, comprehensive investigations of the physico-chemical properties of αSyn condensate remains incomplete. Here, we showed that αSyn condensate possesses a highly negative electrostatic potential that spans the whole condensate. This property causes differential partitioning of dye-labeled αSyn as well as fluorescent molecules by an order of magnitude depending on their net charges. Consistent with this, the phase separation propensity of αSyn is governed by a delicate balance between self-association of αSyn and electrostatic repulsion, hence is antagonized by excess negative charge. We further demonstrated that, in differentiated neuron-like SH-SY5Y cells, αSyn also forms negatively charged condensate. Our results highlighted the significant impact of αSyn condensate’s electrostatic potential on molecular partitioning, hence calling for close examination of the electrostatic property of other biomolecular condensates.

## Introduction

Parkinson’s disease (PD) is the second most prevalent neurodegenerative diseases after Alzheimer’s disease. Despite having a complex etiology, synaptic protein α-synuclein (αSyn) has always been the prime suspect in PD. Notably, αSyn is a major component of Lewy bodies, which are characteristic intracellular inclusion bodies found in the disease-affected brain region of PD patients.^1–2^ αSyn is predominantly structural disordered in its monomeric form,^3–5^ but also exists in multiple oligomeric and fibrillar forms that can be pathogenic.^6–8^ Even though it is widely believed that the aggregation of αSyn plays a critical role in PD, a unified picture of what is going on with αSyn in vivo during the progression of PD remains lacking.^9–10^

Recent studies have shown that αSyn can undergo liquid-liquid phase separation (LLPS),^11^ while the phase-separated droplets maturate over days to form hydrogel that eventually nucleates the formation of amyloid fibrils.^12–13^ Similar mechanism that predisposed a mutated protein to its aggregated form via phase-separated state has also been proposed for FUS^14–15^ and TDP-43^16^ in amyotrophic lateral sclerosis, and tau^17–19^ in Alzheimer’s disease. Hence, it is conceivable that aggregation of these proteins promoted by their LLPS may be a common mechanism that underlies the respective neurodegenerative diseases. Naturally, this insight not only inspires a divergent therapeutic strategy that targets to modify protein LLPS,^20^ while also warrants an in-depth investigation of the molecular partitioning mechanism in these condensates.^16^

αSyn is an acidic protein carrying a predicted net charge of about -9 at pH 7.4. Here, we found that αSyn condensate formed via LLPS possesses a highly negative electrostatic potential such that the dye-labeled αSyn were partitioned differently by as much as ten times depending on the net charge of the dye. The negative electrostatic potential was supported by zeta potential measurement of the condensates. Using cyanine and riboflavin as examples, we further demonstrated that small molecules can be partitioned differently into αSyn condensate based on their charges. Interestingly, we found that the potency of αSyn LLPS is governed by a balance of associative interactions and repulsion between αSyn with excess charge, and the properties of the condensates formed are also dependent on the dye’s charge. In line with the in vitro LLPS experiments, αSyn condensates in the cell were found to be negatively charged. The phenomenon of biomolecular condensates carrying surface electrostatics has been shown recently by two groups.^21–22^ Our work went further to exemplify a tremendous impact of αSyn condensate’s electrostatic potential on molecular partitioning, as well as to provide evidence for such a phenomenon to occur inside the cell.

## Results

### Distribution of dye-labeled αSyn is dependent on charge

To test whether the partitioning of dye-labeled αSyn is affected by electric charge, a group of cyanine fluorophores, i.e. AF647, dsCy5, and Cy5 with the respective net charge of –3, –1, and +1, were chosen to label αSyn(A140C) via thiol-maleimide reaction (Figure 1a). The C-terminal alanine residue was chosen as the site for conjugation in order to minimize disruption to αSyn’s ensemble structures as well as its inter-molecular interactions. As shown by circular dichroism, the secondary structure of native αSyn is largely preserved in all cyanine-labeled αSyn (Supporting Figure S1). We opted to prepare the αSyn LLPS sample for fluorescence microscopy by drop-casting it onto a glass-bottom dish and sealing the coverslip placed on top of it using a UV-LED resin. We found that this enables a consistent assay of αSyn LLPS while greatly alleviating the evaporation problem over a long period of observation (Supporting Figure S2). Furthermore, according to IUPAC recommendation,^23^ we designated the ratio of total concentration of either dye-labeled αSyn or organic fluorophore within the droplets (regardless of its chemical form) to its total concentration outside the droplets as the *distribution ratio*, instead of the commonly used *partition coefficient*, which strictly refers to its un-ionized form.

**Figure 1.**
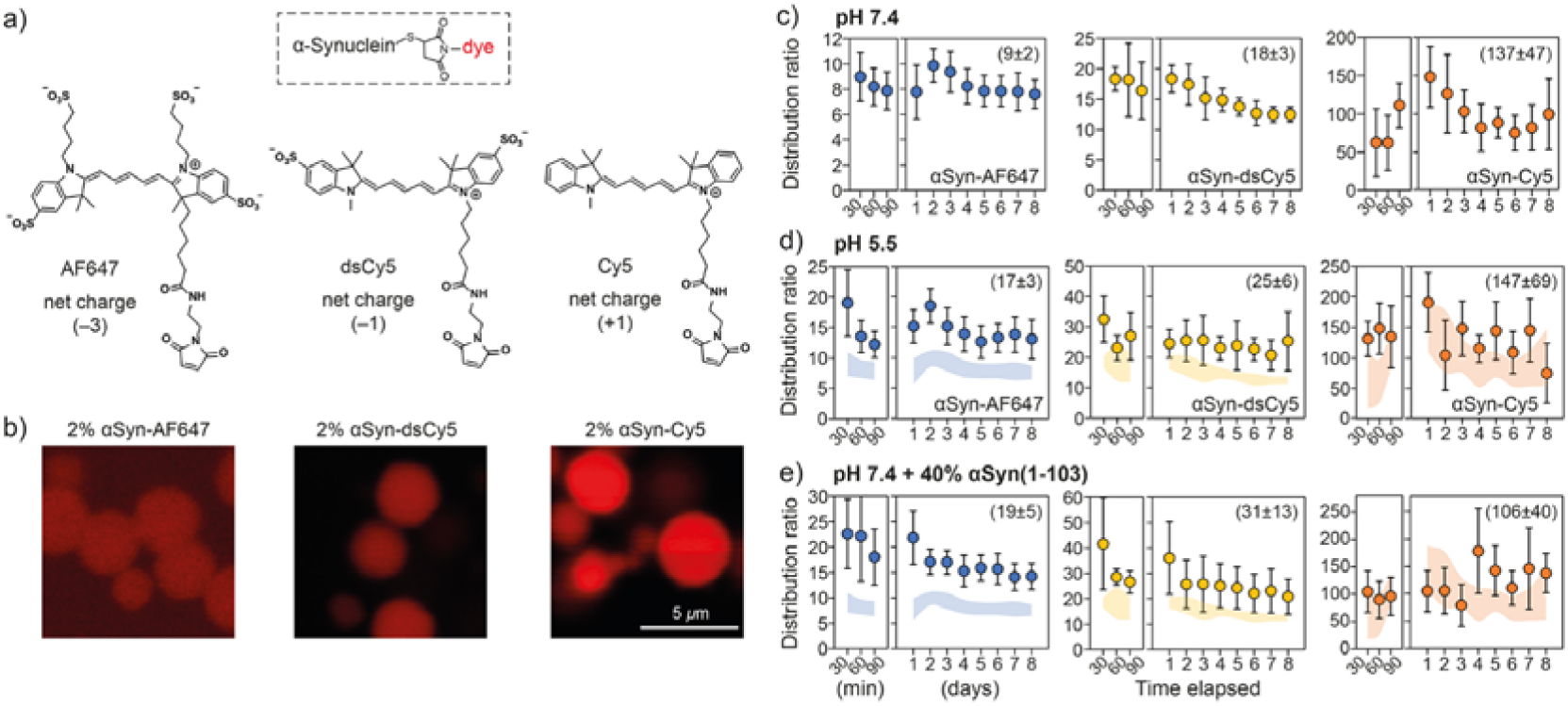
Partitioning of αSyn labeled with tag of different charges. (a) Molecular structures of Alexa Fluor™ 647 C_2_ maleimide (AF647), sulfo-Cy5 maleimide (dsCy5), and Cy5 maleimide (Cy5), with their respective net charges indicated. Michael addition of a thiol group to the maleimide results in a single maleimide–thiol conjugate between αSyn(A140C) and the respective dye. (b) Fluorescence images of condensate formed by 200 μM αSyn doped with 2% molar equivalent of either AF647-, dsCy5-, or Cy5-labeled αSyn at pH 7.4. (c-e) Distribution ratios of AF647-, dsCy5-, and Cy5-labeled αSyn, respectively, at the indicated time points post-LLPS, of samples prepared either in (c) pH 7.4 LLPS buffer, (d) pH 5.5 LLPS buffer, or (e) pH 7.4 LLPS buffer as a mixture of 120 μM αSyn and 80 μM αSyn(1-103). The data represent meanL±Ls.d. of 50-60 data points from two independent samples. Shaded areas in (d) and (e) represent meanL±Ls.d. of the data in (c). The respective average values from d1 and d2 were presented in round bracket.

Sparse, small-sized and highly mobile droplets started to form immediately upon sample preparation, as observed by both fluorescence and differential interference contrast microscopy. The size of the droplets (about 1 to 5 μm) appeared to stabilize after 60 to 90 minutes, and the distribution ratios of the dye-labeled αSyn were quantified up to 8 days post-LLPS. Clearly, the partitioning of the three different cyanine-labeled αSyn were distinct and correlated to the charge of their respective tag (Figures 1b and c), i.e. greater enrichment within the droplets with a more positive tag, thus hinting at the existence of negative electrostatic potential within the droplets. In our case, αSyn-Cy5 was at least ten times more enriched than αSyn-AF647, while αSyn-dsCy5 had double the distribution ratio of αSyn-AF647.

In fact, the distribution ratio can be used to calculate the generalized standard free energy of transfer (Δ*G*^tr^) via the relationship Δ*G*^tr^ = –*RT*ln*K*, where *R* is the gas constant (8.314 J/mol K), *T* is the temperature in Kelvin, and *K* is the distribution ratio.^24–25^ Assuming that the difference in the distribution ratio between αSyn-dsCy5 (relative charge = –1) and αSyn-AF647 (relative charge = –3) can be accounted for solely by the electrostatic repulsion of two additional unit charge, we obtained ΔΔ*G*^tr^ (which is equivalent to *U*_E_ for electric potential energy) of –1.8 kJ/mol at 37 °C. Hence, the electrostatic potential of αSyn condensate was estimated to be –9.3 mV based on *V* = *U*_E_/*nF*, where *n* is unit elementary charge and *F* is Faraday constant (96485 C/mol). Based on the same principle, untagged αSyn (relative charge = 0) and αSyn-Cy5 (relative charge = +1) were expected to have a distribution ratio of 25 and 51, respectively. Since the measured distribution ratio of αSyn-Cy5 was more than 100 (Figure 1c), the cyanine moiety of αSyn-Cy5 is likely to have additional interactions with αSyn condensate beyond the described electrostatic field (see below).

If the electrostatic potential of αSyn condensate was directly proportional to αSyn’s predicted net charge (contributed by its ionizable side chains), then as the protein’s net charge becomes more positive, αSyn-AF647 and αSyn-dsCy5 are expected to become more enriched while αSyn-Cy5 become less. We simulated this situation using two conditions: (i) pH 5.5 buffer solution, and (ii) a mixture of full-length αSyn and a C-terminally truncated αSyn (αSyn(1-103)), in which case the predicted net charges are –6.5 and –4.1, respectively, compared to –9.7 for αSyn at pH 7.4 (Supporting Figure S3). The truncated form of αSyn(1-103) is present physiologically and found in Lewy bodies.^26^ In both conditions, the distribution ratios of αSyn-AF647 and αSyn-dsCy5 were indeed elevated: αSyn-AF647 by 90% and 110% at pH 5.5 and in mixture with αSyn(1-103), respectively, and αSyn-dsCy5 by 40% and 70%, respectively (Figures 1d and e). On the other hand, αSyn-Cy5 also became more enriched at pH 5.5, albeit by about 7% only (Figure 1d) and had a slightly lower distribution ratio in mixture with αSyn(1-103) (Figure 1e). These results indicated that, although the increased distribution ratios of αSyn-AF647 and αSyn-dsCy5 supports the idea of αSyn’s net charge dictating the condensate’s electrostatic potential, the overall effect might include concurrent changes to αSyn’s structural ensembles within the condensate at different experimental conditions. It is known that αSyn becomes structurally more compact and aggregation-prone as the pH decreases.^27–28^

### Negative zeta potential of **α**Syn condensate

Zeta potential is the electrical potential at the slipping plane of the double layer of dispersed particles.^29^ Given the evidence of αSyn condensate possessing negative electrostatic potential that had responded to either pH change or mixing with a truncated αSyn, we attempted to measure zeta potential of the condensates formed at different conditions using a commercial instrument. Since the samples contained a substantial amount of polyethylene glycol (PEG), a single peak with a single-digit negative zeta potential was obtained on the samples without phase-separated αSyn (Figure 2a). We later double-checked on the identity of this peak in sample with either higher concentration of PEG or PEG with higher concentration of NaCl, and still obtained a single-digit zeta potential (Supporting Figure S4).

**Figure 2.**
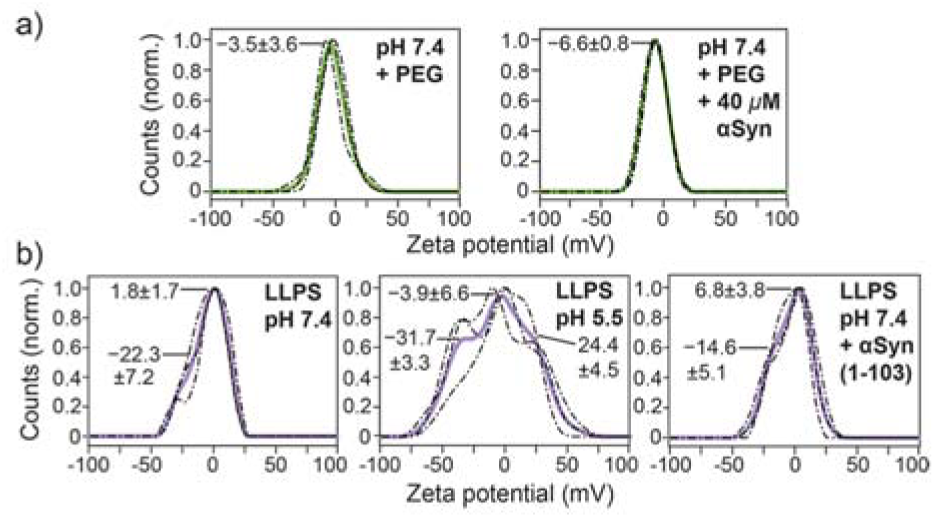
Zeta potential of αSyn condensate. Distribution of the particle’s zeta potential measured on (a) pH 7.4 LLPS buffer with and without 40 μM αSyn and (b) the LLPS samples containing 200 μM αSyn at different conditions, i.e. pH 7.4, pH 5.5, and pH 7.4 as a mixture with 40% αSyn(1-103), respectively. Solid curves were obtained from an average of the triplicate measurements (dashed curves). Zeta potential of the peaks denoted by straight lines were obtained by fitting the distributions to the respective Gaussian mixture models.

When the phase-separated samples were measured, additional peak with a lower amplitude was clearly observed on the left side of the more dominant peak that was attributed to PEG (Figure 2b), hence supporting the presence of negative zeta potential on the surface of αSyn condensate. However, the broad distributions failed to provide quantitative information on any possible differences between the experimental conditions. We noticed that resolution of the distribution was dependent on the maximum voltage applied. Owing to the samples’ high conductivity, the maximum voltage was capped at 50 V during these measurements. In addition, high polydispersity in the zeta potential of biomolecular condensates^21^ could also be a factor. The conductivity of these samples as well as the measured electrophorectic mobility were presented in Supporting Information (Supporting Table S1). Note that an unknown species with a positive zeta potential was also observed in the LLPS sample at pH 5.5, which might be an artefact caused by electrode-analyte reaction.

### Net charge-dependent molecular partitioning into **α**Syn condensate

Since electrostatic potential had a significant effect on the enrichment of cyanine-labeled αSyn, αSyn condensate most probably can modulate the partitioning of chemical compounds based on the same principle. As a proof of principle, we used directly the three cyanine-maleimide fluorophores that are structurally similar except for their net charges, and tested them on the LLPS sample of native αSyn, in which the thiol-maleimide reaction was absent. Being freely diffusing dyes, the cyanine-maleimide were partitioned surprisingly strongly into αSyn condensates, but to different extents depending on their respective net charges (Figures 3a, d). From the respective distribution ratio of AF647 (2.8) and dsCy5 (5.8), we obtained their ΔΔ*G*^tr^ of –1.9 kJ/mol, which is very close to the value calculated for αSyn-AF647 and αSyn-dsCy5. This result supported the conclusion that electrostatic repulsion plays a major role in the differential partitioning of the two cyanine-labeled αSyn. Similar to the case of αSyn-Cy5, Cy5-maleimide had a distribution ratio of 46.6, which is larger than the expected value of 12. This can be explained by the enhanced binding of Cy5-maleimide to αSyn due to the lack of sulfonate groups (Figure 3f).

**Figure 3.**
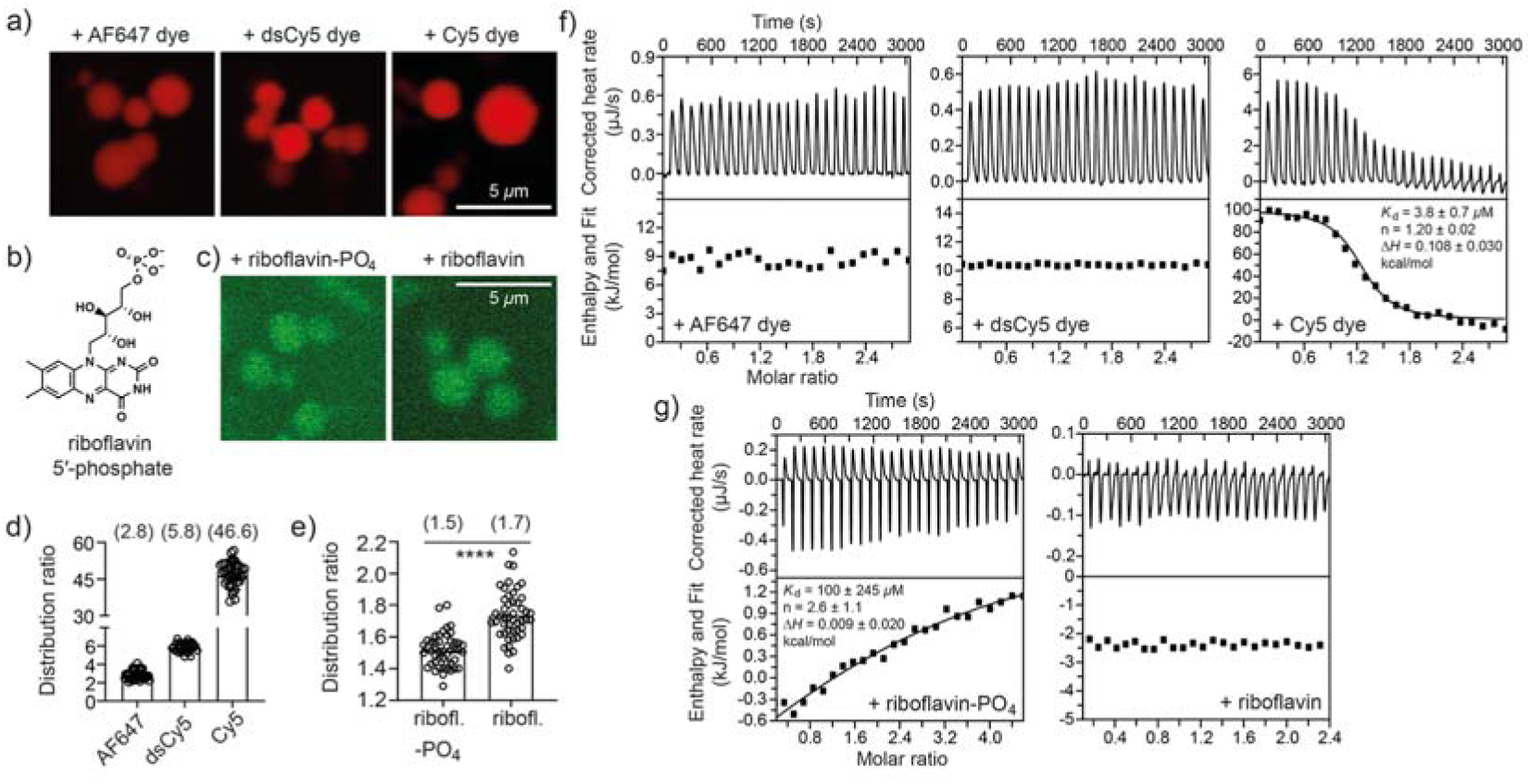
Partitioning of fluorophore with different net charges. (a, c) Fluorescence images of αSyn condensates incubated with (a) 20 μM AF647-, dsCy5-, and Cy5-maleimide, respectively, and with (c) 20 μM riboflavin-PO_4_ and riboflavin, respectively, at pH 7.4 and d1 post-LLPS. (b) Structure of riboflavin-5’-phosphate (riboflavin-PO_4_). (d-e) Distribution ratios of the respective fluorophores determined from the samples in (a, n = 40) and (c, n = 48), with the mean values presented in round bracket. **** indicated P□≤□0.0001 in a two-tailed t-test. (f-g) ITC thermograms and binding isotherms for the titration of the respective fluorophores into αSyn solution.

Likewise, we tested the partitioning of another two fluorophores, riboflavin and riboflavin-5’-phosphate, which differ from each other by the inclusion of a negatively charged phosphate group. Indeed, riboflavin-5’-phosphate exhibited a weaker partitioning compared to riboflavin (Figures 3b-c, e), which reconciled with our overall findings.

Lastly, in order to investigate whether the differential partitioning can be accounted for by the binding affinity of the respective fluorophores to αSyn protein, we ran the isothermal titration calorimetry (ITC) experiments by titrating the fluorophores into αSyn solution (Figures 3f-g). Both AF647- and dsCy5-maleimide exhibited negligible binding to αSyn, while Cy5-maleimide bound to αSyn with *K*_d_ of 3.8 ± 0.7 μM, indicating that specific binding of Cy5-maleimide might contribute to its large distribution ratio in phase-separated αSyn (Figure 3f). On the other hand, riboflavin-5’-phosphate bound to αSyn with *K*_d_ of 100 ± 245 μM, while riboflavin had negligible binding to αSyn (Figure 3g). Even though riboflavin-5’-phosphate has a higher affinity to αSyn than riboflavin, it was less enriched in αSyn condensate. In overall, these results indicated a strong correlation between the fluorophores’ net charge and their partitioning into αSyn condensate. This correlation is due to their interaction with the negative electrostatic potential within αSyn condensate, apart from the partial contribution of their binding to αSyn protein.

### αSyn LLPS is antagonized by excess charge

The homogeneous fluorescence intensity of dye-labeled αSyn (Figure 1b) and freely diffusing dyes (Figures 3a, c) within the interior of the condensate’s cross section hinted at a rather uniform distribution of charges. We hypothesized that LLPS of αSyn might be dictated by a delicate balance between self-association of αSyn and repulsion of negative charges that were not screened, hence generating condensate with uniform charge density on the condensate’s length scale. To test it, we prepared LLPS samples with increasing proportion of dsCy5-labeled αSyn, simulating the addition of excess negative charges. Indeed, the number of droplets dropped drastically as the percentage of αSyn-dsCy5 increased (Figure 4a). With 100% αSyn-dsCy5, no droplets were observed after 90 minutes, and after one day of incubation, very sparsely distributed droplets could be found accompanied by irregular-shaped aggregates. This result showed that, under our experimental condition, LLPS of αSyn was antagonized by excess negative charge.

**Figure 4.**
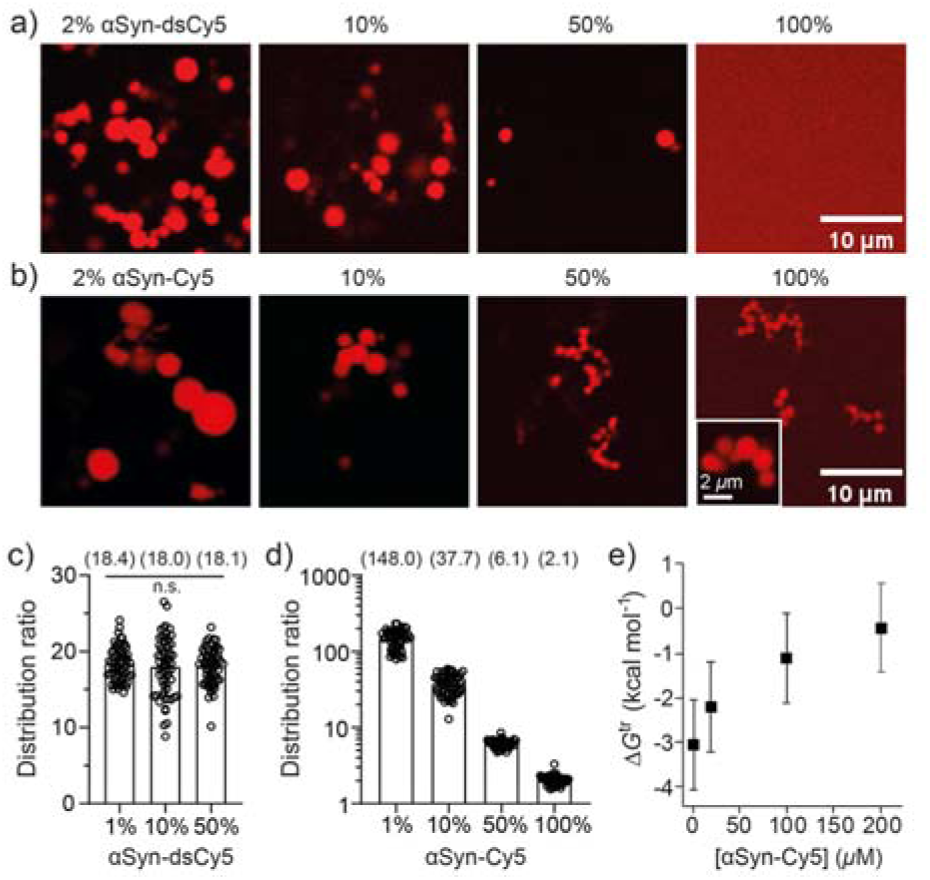
Electrostatic effect on αSyn LLPS propensity. (a-b) Fluorescence images of αSyn LLPS samples prepared by mixing different proportions of untagged αSyn and either (a) αSyn-dsCy5 or (b) αSyn-Cy5 at pH 7.4 and d1 post-LLPS. Distribution ratios of (c) αSyn-dsCy5 and (d) αSyn-Cy5, respectively, in the LLPS samples containing different proportions of untagged and dye-labeled αSyn (n=60 from duplicate samples). Mean values were presented in round bracket. Welch and Brown-Forsythe ANOVA tests were performed on (c). (e) Mean Δ*G*^tr^ of αSyn-Cy5 calculated from the data in (d). Error bar represents standard deviation.

We then performed the same type of experiment using αSyn-Cy5, which carries an additional positive charge instead. In this case, formation of round-shaped droplets could be observed throughout the samples up to 100% of αSyn-Cy5, even though there was a clear trend of decreasing sizes (Figure 4b). Apparently, the property of the condensate had been modified by the addition of αSyn-Cy5, as shown by the decreasing distribution ratio (Figure 4d). This was in contrast to the experiment with increasing proportion of αSyn-dsCy5, in which distribution ratio remained essentially constant (Figure 4c).

When Δ*G*^tr^ remained constant as the proportion of αSyn-dsCy5 was increased, it implies either LLPS is driven equally by αSyn:αSyn (homotypic) and αSyn:αSyn-dsCy5 (heterotypic) interactions^24^, or αSyn-dsCy5 does not support LLPS at all. The absence of LLPS at 100% αSyn-dsCy5 showed that αSyn-dsCy5 behaved like a “client”^30^ in the condensate formed by untagged αSyn, thus αSyn-dsCy5 would have minimal effect on the phase-separated state of untagged αSyn. On the other hand, αSyn-Cy5 acted as the “scaffold”^30^ together with untagged αSyn to form condensates with variable distribution ratios depending on the ratio of their concentrations. Its Δ*G*^tr^ exhibited a curve similar to the results on intracellular NPM1 condensate,^24^ showing that heterotypic αSyn:αSyn-Cy5 interaction is more stabilizing, and charge neutralization may play a role in their LLPS (Figure 4e). These results indicated that LLPS propensity and the resulting phase-separated states of αSyn are highly sensitive to the mixture composition of αSyn modified by moiety of distinct net charges.

### Negatively charged **α**Syn condensate in the cell

We went on to study LLPS of αSyn in the cell using different cell lines, some of which had been reported by other groups with disparate observations.^12, 31–33^ Firstly, in COS-7 cells, neither the overexpression of αSyn alone nor its co-expression with Synapsin-1 resulted in any observable αSyn droplets (data not shown). Secondly, in HEK293T cells, only a very small percentage of co-transfected cells exhibited αSyn droplet formation (Supporting Figure S5). Conversely, a robust αSyn LLPS was achievable by overexpression of αSyn-mCherry in differentiated SH-SY5Y cells – a model of cholinergic/dopaminergic neuron for PD research^34^ (Figure 5a). Notably, these αSyn droplets exhibited a spherical shape with a uniform size of about 0.5 μm and were dispersed throughout the cytosol, unlike the observation of larger-diameter, perinuclear aggresome as reported by others.^32^ Time-lapse imaging showed that these αSyn condensates in SH-SY5Y cell were highly mobile. Upon analyzing the movement of these condensates, multiple incidents of collided condensates that resisted fusion had been observed (Figure 5b and Supporting Videos V1-3), thus hinting at the existence of surface electrostatics on these condensates based on the same principle demonstrated in vitro.^21^

**Figure 5.**
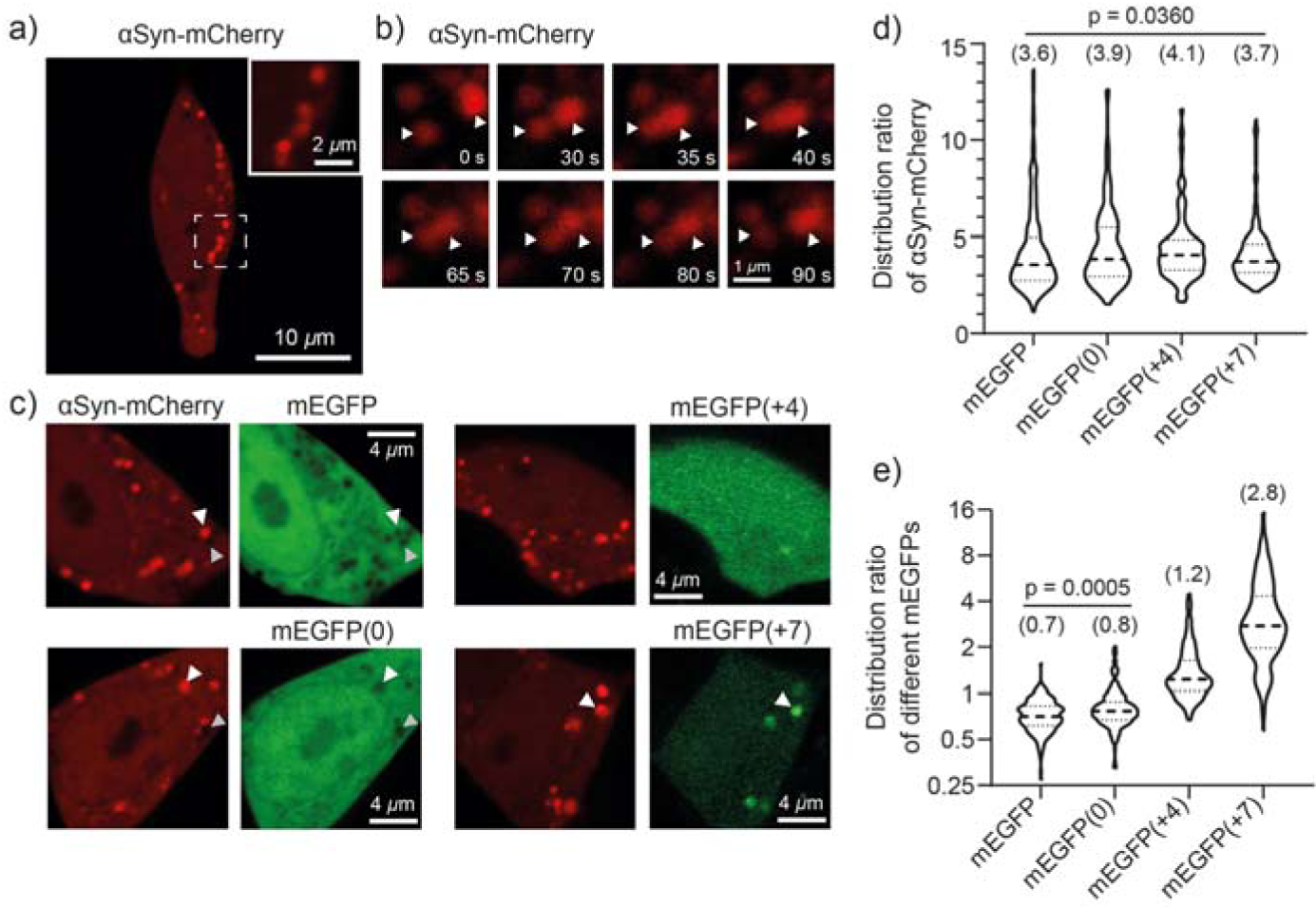
Negatively charged αSyn condensate in the cell. (a) Representative fluorescence image of SH-SY5Y cells overexpressing αSyn-mCherry. (b) Snapshots illustrating the 90-second dynamic movement of two αSyn-mCherry droplets that resisted fusion (denoted by white arrowhead). (c) Fluorescence images of SH-SY5Y cells co-expressing αSyn-mCherry and the respective wild-type or mutant mEGFP carrying distinct net charge indicated in bracket. For wild-type mEGFP and mEGFP(0), examples of mEGFP excluded from αSyn condensate were denoted by white arrowheads, while cavities that excluded both αSyn-mCherry and mEGFP were denoted by grey arrowheads. For mEGFP(+7), an example of enriched mEGFP was denoted by white arrowheads. Violin plots of the distribution ratio of (d) αSyn-mCherry and (e) different mEGFP(X) determined from the respective combination of co-expressed cells as shown in (c). For each combination, n=230 from 20 cells in duplicate cultures. Median (indicated in bracket) and quartiles were denoted by dashed and dotted lines. Kruskal-Wallis test was performed on data in (d), while Mann-Whitney test was performed on mEGFP and mEGFP(0) in (e).

In order to further determine whether a negative electrostatic potential does occur within the αSyn condensates, we co-transfected αSyn-mCherry together with different mEGFP possessing distinct net charge in SH-SY5Y cells (Figure 5c), and measured their distribution ratios akin to the in vitro experiments. Wild-type mEGFP has a net charge of -8, while three other mutant mEGFPs with a net charge of 0, +4, and +7, respectively, were generated based on the protein supercharging principle^35^ (refer to Supporting Figure S6 for amino acid sequences). Clearly, αSyn condensates appeared to partially exclude mEGFP and mEGFP(0), while mEGFP(+4) was distributed homogenously and mEGFP(+7) was enriched in the condensates (Figure 5c). Quantification of their distribution ratios showed that mEGFP(+7) was enriched by about 2.8 times, and both wild-type mEGFP and mEGFP(0) had a distribution ratio of less than 1 while mEGFP(0)’s was slightly larger than that of wild-type mEGFP (Figure 5e). Overall, the trend showed a correlation between mEGFP’s net charge and its partitioning, hence strongly supporting the existence of a negative electrostatic potential in αSyn condensates. In all cases, αSyn-mCherry exhibited only slight variation in distribution ratio (Figure 5d), indicating that overexpressed mEGFP(X) acted as a “client” to αSyn condensate. Another interesting observation to take note was the cavities that appeared to exclude both αSyn-mCherry and mEGFP (Figure 5c, grey arrowheads), which indicated a possible existence of other negatively charged condensates.

## Discussion

In this work, we embarked on a careful analysis of the distribution ratio of αSyn labeled with tags of distinct net charges, and identified certain pitfalls that may be encountered in biomolecular LLPS research. Firstly, our data pointed out the caveat of utilizing a charged fluorescent tag for quantifying the distribution ratio of a biomolecule upon LLPS, particularly for charged condensates. As shown, αSyn condensate is a negatively charged simple coacervate, while FUS (simple coacervate) and PR_25_:PolyU (complex coacervate) were also reported to carry negative zeta potentials,^21^ thus we anticipated more biomolecular condensates to possess electrostatic potential.

Secondly, our results also highlighted the impact of the tag’s net charge on the thermodynamics of αSyn LLPS, namely whether LLPS can happen and how the phase-separated states differ depending on the mixture composition of tagged αSyn. When labeled with tags of opposite net charge, αSyn became either a “client” or a “scaffold” in the phase-separated states, demonstrating a complex interplay of many factors in which electrostatics played a major role. For the samples containing αSyn-dsCy5 only, LLPS was inhibited, indicating a surplus of repulsive interactions that had overcome the *capacity* of αSyn condensate in that condition, thereby altering the system’s Gibbs free energy of mixing.^36^ We speculated that this mechanism might explained why LLPS of tetracysteine-tagged αSyn in HeLa cells was substantially enhanced only after incubation with either ammonium ferric citrate or copper sulphate,^12^ in which case, excess negative charge on αSyn could be partially neutralized by the cations to promote its LLPS.

Molecular partitioning is an important aspect in the study of biomolecular condensates, since local segregation of specific biomolecules is a putative function of LLPS in the biological system.^37–38^ *Aberrant phase separation* generally refers to a cellular state in which the regulation of LLPS of a biomolecule has gone haywire causing a disease,^39–40^ though most of the examples focused on the sol-gel phase transition of condensates that eventually produced various types of pathogenic aggregates^41^. Until recently, there is an increased interest and reports on the disease implication of *aberrant partitioning* of biomolecules, which were also electrostatically driven.^42–43^ For example, a frameshift mutation renders the nuclear protein HMGB1 more positively charged and being mispartitioned into nucleolus, which then causes a rare complex malformation disease.^42^ Furthermore, based on the analysis of the variants of other disease-related genes, the mechanism of nucleolar mispartitioning and dysfunction could even be linked to many other genetic diseases.^42^ We have provided evidence that αSyn-mCherry forms negatively charged condensate in differentiated SH-SY5Y cells, thereby presenting a useful cellular model for the investigation of the physiological consequence of αSyn LLPS and its relationship with PD beyond amyloid fibril formation.

The partition coefficient of various drug molecules is routinely used by pharmaceutical industry as a measure of the molecule’s hydrophobicity and a proxy for its membrane permeability.^44^ However, taking aside the problem of drug delivery to the cell, successful drug targeting with a positive pharmaceutical outcome still requires many other considerations, particularly accumulation of the drug at the right location within the cells. This concept was nicely demonstrated by a recent report on the differential partitioning of certain cancer drugs into distinct biomolecular condensates that subsequently demarcates their therapeutic efficacy.^45^ Our results showed that electrostatic potential in the condensate can exert a huge effect on the partitioning of small molecules. In the case of αSyn condensate, enrichment of Cy5-maleimide was ten times greater than dsCy5-maleimide, given that they are structurally similar except for a charge difference of 2*e*. Although specific binding of Cy5-maleimide to αSyn does contribute a huge part to the partitioning, enrichment of Cy5-maleimide is expected to be double that of dsCy5-maleimide solely due to electrostatic interaction. Nevertheless, developing a general model that relates the extent of enrichment to the net charge of a molecule, as well as finding out whether the molecule-protein binding equilibrium could be altered within a charged condensate, would require further studies.

In conclusion, we have demonstrated that αSyn condensate, formed either in vitro or in the cell, carries negative electrostatic potential that spans the cross section of the condensate, hence is able to modulate the partition of molecules according to their net charges. In addition, net charge of the fluorescent tag exerted a significant impact on αSyn LLPS, highlighting the importance to re-examine the electrostatic potential of various biomolecular condensates and its effect on LLPS propensity and molecular partitioning.

## Materials and Methods

### Materials

Cyanine dyes were purchased from the respective companies: Alexa Fluor™ 647 C2 maleimide (Catalog No. A20347, Thermo Fisher Scientific), diSulfo-Cy5 maleimide (Catalog No. BDC-47, CAS No. 2130955-10-9, CONFLUORE), and Cy5 maleimide non-sulfonated (Catalog No. A8139, APExBIO). Riboflavin (CAS No. 83-88-5), riboflavin-5’-phosphate sodium salt dehydrate (riboflavin-PO4, CAS No. 6184-17-4), NaOH (CAS No. 1310-73-2), Tris-HCl (CAS No. 1185-53-1), NaCl (CAS No. 7647-14-5), MES (CAS No. 145224-94-8) and EDTA (CAS No. 60-00-4) were all purchased from Sigma. PEG-8000 (CAS No. 25322-68-3) was purchased from Solarbio. IPTG (Catalog No. ST098, CAS No. 367-93-1), DTT (Catalog No. ST043, CAS No. 3483-12-3), PMSF (Catalog No. ST505, CAS No. 329-98-6), HEPES (Catalog No. ST092, CAS No. 7365-45-9) and TCEP (Catalog No. ST045, CAS No. 51805-45-9) were all purchased from Beyotime. The reagents used for cell culture and transfection were from Beyotime, Gibco, and Thermo Fisher Scientific, unless otherwise specified, including BDNF (Brain Derived Neurotrophic Factor, Catalog No. 450-02-10, PEPROTECH) and EC23 (Catalog No. HY-12309 EC23, CAS No. 2130955-10-9, MedChemEpress). All solutions were freshly prepared and filtered through a 0.22 μm syringe filter for each use.

## Protein expression and purification

Expression and purification of human full-length αSyn, αSyn(A140C) and αSyn(1-103) were performed by a combination of purification steps.^46^ Briefly, colonies of Escherichia coli BL21(DE3) transformed with the respective expression vectors were cultivated in LB broth (100 mg/L ampicillin) at 37 °C with shaking (220 rpm). At A600 of 0.6-0.8, the cultures were induced with 1 mM isopropyl-β-D-thiogalactoside (IPTG) for 4 h. The cells were then harvested by centrifugation (10000 rpm, 10 min) and the pellet was kept frozen at -80 °C. For protein purification, the pellet was resuspended in lysis buffer (10 mM Tris-HCl, pH 8.0, 1 mM EDTA, 1 mM DTT, 1 mM PMSF) and sonicated at 300 W for 20 min. Cell debris was then separated by centrifugation (10000 rpm, 20 min), and the supernatant was added 1.5% HCl to pH 3.5, followed by centrifugation (10000 rpm, 10 min) to remove unwanted proteins. The supernatant was added 1 M NaOH to pH 7.5. Then, 8.7 g ammonium sulfate was added for αSyn to precipitate at 4 °C for at least 1 hour. Subsequently, the solution was centrifuged at 10000 rpm and 4 °C for 20 min to pellet down αSyn. The pellet was solubilized and dialyzed against the buffer (20 mM Tris-HCl, pH 8.0, 1 mM EDTA, 150 mM NaCl) overnight at 4 °C using a 10 kDa MWCO mini-dialysis unit (Catalog No. MD34-10KD, RuiTaibio). This process was carried out to remove salts. Finally, the protein solution was loaded onto HiLoad 26/600 Superdex 200 pg column attached to an AKTA purifier (GE Healthcare) and eluted isocratically at 4 °C in the buffer (25 mM HEPES, pH 7.4, 150 mM NaCl). The monomeric αSyn fraction was collected and lyophilized to powder. Lyophilized αSyn was dissolved in two different buffers: (i) 25 mM HEPES, 150 mM NaCl, pH 7.4; or (ii) 25 mM MES, 150 mM NaCl, pH 5.5, and then aliquoted, flash frozen and stored at -80°C for future use. Purity of the protein was confirmed by running sodium dodecyl sulfate-polyamide gel electrophoresis (SDS-PAGE, Beyotime).

### Fluorophore labeling of **α**Syn

The maleimide dyes (AF647, dsCy5 and Cy5) were dissolved in DMSO at the concentration of 10 mM. Eight to ten-fold molar excess of the individual dye was added to 100 μM αSyn(A140C) protein (TCEP pre-treated for 30 min), and incubated at room temperature for 1 h. The reaction mixture was then diluted with 1 mL buffer (25 mM HEPES, pH 7.4, 150 mM NaCl), and dialyzed against the same buffer overnight at 4 °C using a 10 kDa MWCO mini-dialysis unit (Catalog No. MD34-10KD, RuiTaibio) to remove unreacted dye. The extent of modification of the protein with the dye was determined by NanoDrop One^C^ (Thermo Fisher) and SDS-PAGE. For another experiment condition (25 mM MES, pH 5.5, 150 mM NaCl), the dye-labeled αSyn(A140C) protein was dialyzed against the target buffer overnight at 4 °C and concentrated by 3000 MWCO Protein Concentrators PES (Catalog No. VS2021, Sartorius). All labeled proteins were confirmed by mass spectrometry (Supporting Figures S7-S11).

### Mass spectrometry

HRMS (ESI) spectra were obtained using a Thermo Fisher Q-Exactive Mass Spectrometer. 0.1 mg/mL proteins were measured in 25 mM HEPES, pH 7.4, 150 mM NaCl (Supporting Figures S7-S11).

### Circular dichroism spectroscopy

30 μM αSyn, AF647-, dsCy5-, and Cy5-labeled αSyn at 50 mM sodium phosphate buffer, pH 7.4 were performed via Applied Photophysics Chirascan circular dichroism spectrometer. Spectra were recorded from 260 to 200 nm at 25 °C with a speed of 100 nm/min via a circular dichroism micro cuvette of 1 cm path length.

### In vitro LLPS assay of **α**Syn

The protein solution was thawed and pre-cleared by centrifugation (10000 rpm, 10 min) at 4 °C, and its final concentration was determined. The LLPS sample was prepared by mixing the components accordingly, with 10% (w/v) PEG-8000 being added at the end and mixed gently to ensure homogeneity. 10 μL of the mixture was drop-casted in the middle of a 35-mm glass bottom microwell dish (Catalog No. P35GC-1.0-14-C, MatTek Corporation), air-exposed on the surface of heat plate (37 °C for 5 min), and then covered with a coverslip and sealed with UV-LED resin (see Figure S2). The dish was then kept at 37 °C for the whole duration of our experiment including microscopy.

### Confocal microscopy

Fluorescence images were acquired on a Zeiss LSM 900 inverted confocal microscope (Carl Zeiss AG) with a Plan-Apochromat 63x/1.40 Oil DIC M27 objective lens. The sample dishes were kept inside a temperature-regulated chamber (37 °C, 5% CO_2_, STXG-WSKMX-SET, Tokai Hit Co., Ltd.) mounted on the microscope stage. Supply of CO_2_ was not used for in vitro LLPS samples. Cyanine dyes were excited by 653-nm laser and detected at 656-700 nm, riboflavin/riboflavin-PO4 were excited by 488-nm laser and detected at 410-561 nm, mEGFP/EGFP were excited by 488-nm laser and detected at 400-560 nm, mCherry was excited by 561-nm laser and detected at 576-700 nm. Laser power was adjusted to avoid saturation of fluorescence intensity. For in vitro LLPS experiment, ten images (20.28×20.28 μm, 512×512 pixels, 16 bit-depth) were acquired for each time point for each sample. For imaging the SH-SY5Y cells, 1 μm-separated z-stack of images (pixel size=0.085×0.085 μm, 16 bit-depth) were acquired for each cell. All images were analyzed using ImageJ software (National Institutes of Health). To determine the distribution ratio, which is the ratio of the mean fluorescence intensity between inside and outside of the droplet, pairs of region of interest (ROI, 0.79 μm diamater circle) were chosen such that one was centered in the droplet and the other was in the vicinity outside the droplet. The brighter droplets were chosen in order to obtain intensity closer to the center of the droplet. For the distribution ratio in SH-SY5Y cells, circular ROIs with 0.34 μm diameter were chosen.

### Zeta potential

Measurement of zeta potential was performed using Zetasizer Nano ZS (Malvern Instruments Ltd.) with disposable folded capillary cell (DTS1070, Malvern Panalytical Ltd.) and analyzed using Zetasizer Software (Malvern Panalytical Ltd.). To acquire large volume of LLPS samples for the measurement, 1.5 mL solution of 200 μM αSyn containing 10% (w/v) PEG-8000 was drop-casted onto a 35-mm glass bottom microwell dish and incubated for 30 min at 37 °C, aspirated and placed inside an Eppendorf tube for incubation of another 2 h at 37 °C. The final volume of the solution was about 1.3 mL, and droplet formation was confirmed by confocal microscopy before zeta potential measurement. Zeta potential was measured at 37 °C three times for each sample. To calculate zeta potential, the parameters for water as the dispersant at 37 °C were used, i.e. viscosity=0.6864 cP and dielectric constant = 74.4. The output zeta potential distribution was interpolated and fitted to different Gaussian mixture models using MATLAB (The MathWorks, Inc.).

### Isothermal titration calorimetry (ITC)

The thermodynamic parameters of the fluorophores binding to αSyn protein were measured using a Nano ITC200 instrument from TA Instruments. Experiments were carried out at 25 °C in buffer (25 mM HEPES, pH 7.4, 150 mM NaCl) with 10% (v/v) DMSO. All measurements were performed two times and all solutions were accurately degassed before the ITC experiments to avoid the formation of air bubbles in the calorimeter vessel. The concentration of AF647-, dsCy5-, Cy5-maleimide and riboflavin was 1 mM, and titrated into αSyn protein solution of 0.1 mM. The concentration of riboflavin-5’-phosphate was 0.5 mM, and titrated into αSyn protein solution of 0.02 mM. 25 injections of 2 μL fluorophore solution were added into a 200-μL cell containing αSyn protein solution, every 250 s. All the measurements were conducted at a continuous stirring rate of 350 rpm. Control experiments were carried out to calculate the heat of dilution. Integrated heat data obtained from the titrations were fitted to the independent model using NanoAnalyze software (TA Instruments) as well as a ITC Data Fitting Program written by Debashish Sahu (https://sites.lsa.umich.edu/debsahu/2016/04/27/itc-data-fitting-program/).

### Growth and differentiation of SH-SY5Y cells

Differentiation of SH-SY5Y cells was performed according to Forster *et al.*^47^ with minor modifications. Cells were maintained at 37 °C and 5% CO_2_ in Dulbecco’s Modified Eagle’s Medium (DMEM, Gibco) supplemented with 10% Fetal Bovine Serum (FBS, Thermo Fisher) and 1% penicillin/streptomycin (BI). T25 cell culture flasks (Sarstedt) were used for cell proliferation. After cells had reached 80-90% confluence, splitting was done by washing cells with phosphate-buffered saline (PBS, Beyotime) once and incubating them with trypsin-EDTA for 2 min at 37 °C. The cell pellet was then resuspended in DMEM medium (10% FBS, 1% pen/strep) and plated in µ-Slide 4 Well Chamber Slide (Catalog No. 80426, tissue culture-treated, ibidi GmbH). After 24 h, culture medium was switched to DMEM/F12 (5% FBS) supplemented with 10 μM all-trans-retinoic acid (EC23) to promote differentiation and induce neuronal phenotype. Three days later, culture medium was switched to Neurobasal medium (containing B27 supplement and GlutaMAX) supplemented with 50 ng/mL of brain derived neurotrophic factor (BDNF). Afterwards, culture medium was replaced every 2-3 days using the same recipe to maintain the differentiated cells. Successful differentiation of SH-SY5Y cells was monitored by morphological assessment of neurite outgrowth. Fluorescent protein (mEGFP or mCherry) was fused to the C-terminus of human αSyn following four repeats of glycine and serine, i.e. αSyn-(GS)4-FP. The respective DNA constructs were cloned into pcDNA3.1(+) vector with a CMV promoter. Transfection of SH-SY5Y cells with the respective αSyn expression vectors was performed using Lipofectamine 6000 (Thermo Fischer) following manufacturer’s instructions. Briefly, DNA (2.4 μg in total) was incubated with Lipofectamine 6000 (5 μL) and Opti-MEM (120 μL) for 10 min at room temperature, and then added into the cells with ∼700 μL culture medium. The cells were incubated for 4-6 h (37 °C and 5% CO_2_), and the culture medium was then replaced by fresh medium.

## Acknowledgement

We are grateful for the generous donation of αSyn bacterial expression vector by Guohua Lv, SH-SY5Y cell line by Keqiang Ye, and the DNA of mouse Synapsin-1 by Mingjie Zhang. We thank Xi Liu for discussion and technical assistance. This work was supported by National Key R&D Program of China (Grant no. 2022YFA1303100, headed by Xuebiao Yao). L.E.W. acknowledged the support of National Natural Science Foundation of China (Grant no. 32350012 to L.E.W.). S.C. and Z.L. acknowledged the support of Shenzhen Science, Technology and Innovation Commission (Grant no. JCYJ20210324140212034 to S.C. and Grant no. KQTD20210811090117032 to Z.L.). Additionally, Z.L. acknowledged the support of Shenzhen Technological Research Center for Primate Translational Medicine (Grant no. XMHT20220104005).

## Author contributions

L.E.W. conceptualized and supervised the project. Q.Y. performed all in vitro experiments and analyzed the data. S.-F.C. performed the cellular experiments and analyzed the data. P.Z. assisted in zeta potential measurements. Z.L. provided resources for the experiments. S.C. provided methodology for the cellular experiments. L.E.W. wrote the manuscript. All authors commented on the manuscript.

## Data availability

Original data are available from the corresponding author (L.E.W.) upon request.

