## Supporting Figure for "Negatively charged α-Synuclein condensate modulates partitioning of molecules"

### Supporting Figures:

#### Circular dichroism spectra of native and cyanine-labeled αSyn

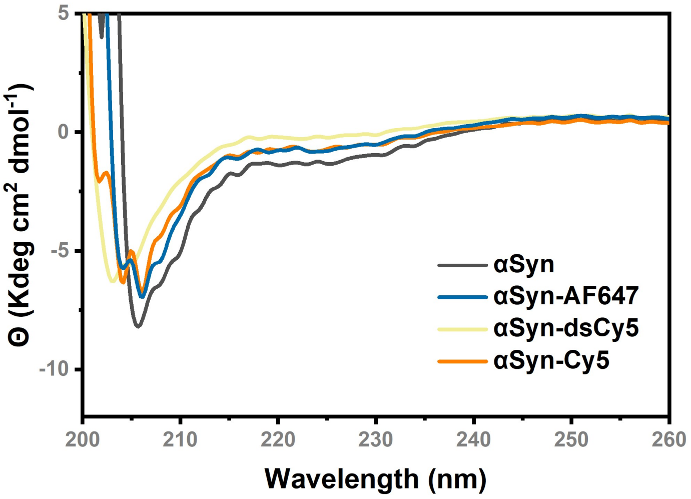

**Figure S1.** CD spectra of 30 $\mu$M αSyn, AF647-, dsCy5-, and Cy5-labeled αSyn at 50 mM sodium phosphate buffer, pH 7.4.

**Preparation of αSyn LLPS samples**

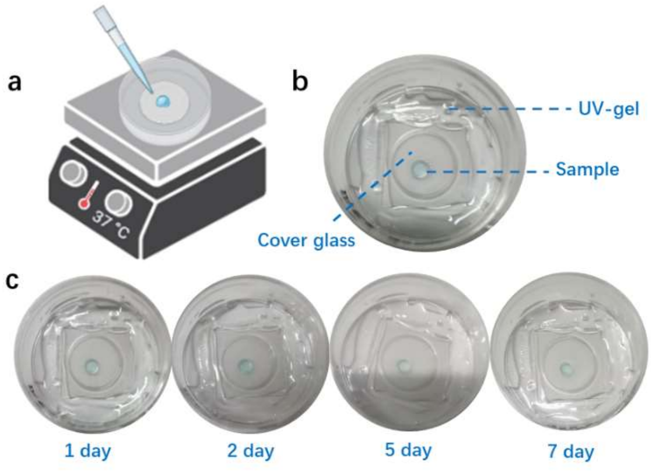

**Figure S2.** (a) Schematic depicting the preparation of αSyn LLPS sample. (b) Photo of a αSyn LLPS sample sealed in the 35-mm glass bottom dish immediately after preparation. (c) A series of photos were taken on the αSyn LLPS sample shown in (b) throughout the duration of the experiment from day 1 to 7, illustrating controlled volatilization and negligible shrinkage of the droplet.

**Net charge of αSyn as a function of pH**

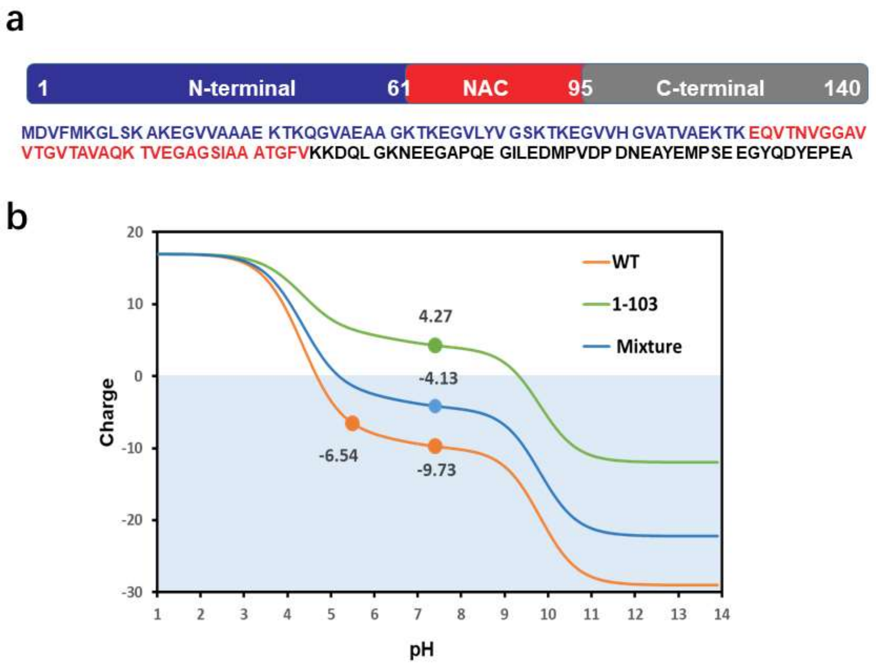

**Figure S3.** (a) Domain architecture of αSyn. The N-terminus (blue), non-amyloid-β component (NAC) (red) and C-terminus (grey) are shown. (b) The net charge as a function of pH, as predicted by Prot pi (https://www.protpi.ch/Calculator/ProteinTool), were plotted for full-length αSyn (WT, orange line), truncated αSyn(1-103) (1-103, green line), and a mixture of full-length αSyn and αSyn(1-103) at the molar ratio of [WT]:[1-103] = 0.6:0.4 (Mixture, blue line).

### Zeta potential of PEG

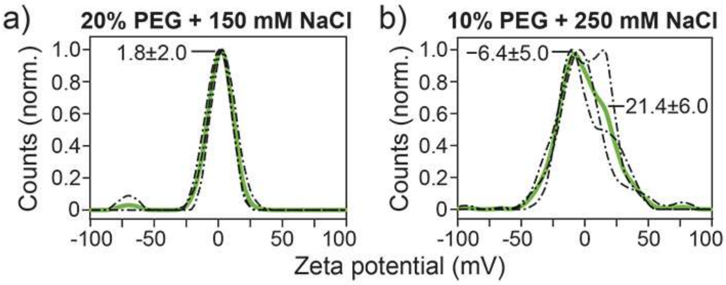

**Figure S4.** Distribution of the zeta potential measured on (a) 20% PEG with 25 mM HEPES, pH 7.4, 150 mM NaCl and (b) 10% PEG with 25 mM HEPES, pH 7.4, 250 mM NaCl.

**Table S1.** Suspension conductivity and electrophoretic mobility values of the zeta potential measurements.

| Conductivity (mS/cm) | | | |
|---|---|---|---|
| 10% PEG | 14.8 ± 0.9 | | |
| 10% PEG + 40 $\mu$M αSyn | 15.9 ± 0.7 | | |
| LLPS pH 7.4 | 24.0 ± 1.3 | | |
| LLPS pH 5.5 | 27.0 ± 0.5 | | |
| LLPS pH 7.4 + αSyn(1-103) | 20.3 ± 1.3 | | |
| Electrophoretic mobility ($\mu$m·cm/V·s) | | | |
| 10% PEG | −0.3 ± 0.4 | | |
| 10% PEG + 40 $\mu$M αSyn | −0.6 ± 0.1 | | |
| LLPS pH 7.4 | −2.3 ± 0.5 | −0.03 ± 0.19 | |
| LLPS pH 5.5 | −3.8 ± 0.3 | 0.3 ± 0.2 | 3.3 ± 2.2 |
| LLPS pH 7.4 + αSyn(1-103) | −1.4 ± 0.5 | 0.7 ± 0.4 | |

**Co-expression of αSyn and Synapsin-1 in HEK293T cells**

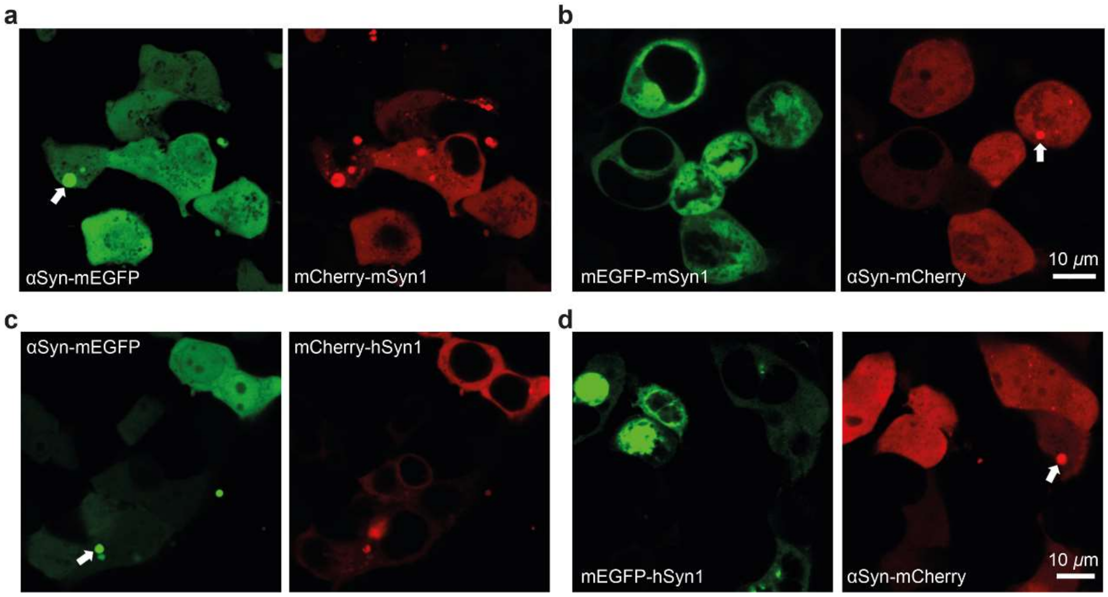

**Figure S5.** Representative fluorescence images of HEK293T cells co-expressing (a) αSyn-mEGFP and mCherry-mSyn1, (b) αSyn-mCherry and mEGFP-mSyn1, (c) αSyn-mEGFP and mCherry-hSyn1, and (d) αSyn-mCherry and mEGFP-hSyn1. Either mEGFP or mCherry was fused to the N-terminus of mouse Synapsin-1 and human Synapsin-1 with a linker of 10 and 14 amino acids, respectively, i.e. FP-LEVLFQGPGS-mSyn1 and FP-LEVLFQGPGSKLAT-hSyn1. All DNA constructs were cloned into pcDNA3.1(+) vector with a CMV promoter.

### Amino acid sequences of mEGFP(X)

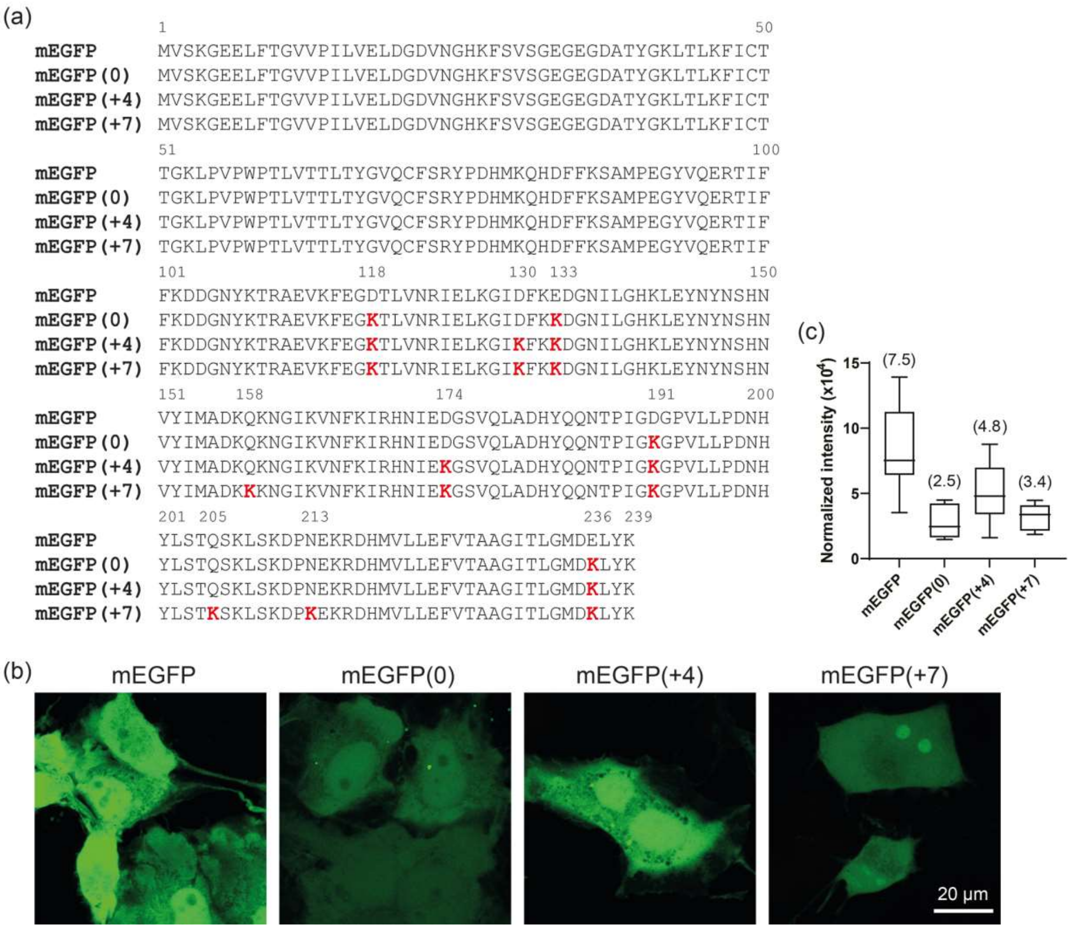

**Figure S6.** (a) Multiple sequence alignment of the members of wild-type mEGFP and three other mutant mEGFPs with a net charge of 0, +4, and +7, respectively. Residues in the mutant mEGFP that are different from wild-type mEGFP are colored in red. (b) Fluorescence images of COS7 cells overexpressing wild-type mEGFP and three mutant mEGFPs. (c) Box and whisker plot of the normalized fluorescence intensity determined from the respective cells in (b). Median is indicated in bracket.

**Mass spectrometry**

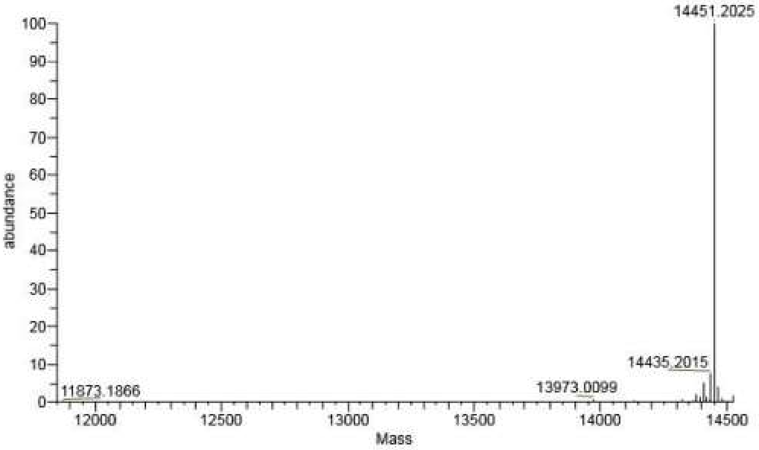

**Figure S7.** The ESI-MS spectrum of WT αSyn. Theoretical mass 14460.16, experimental mass 14451.20.

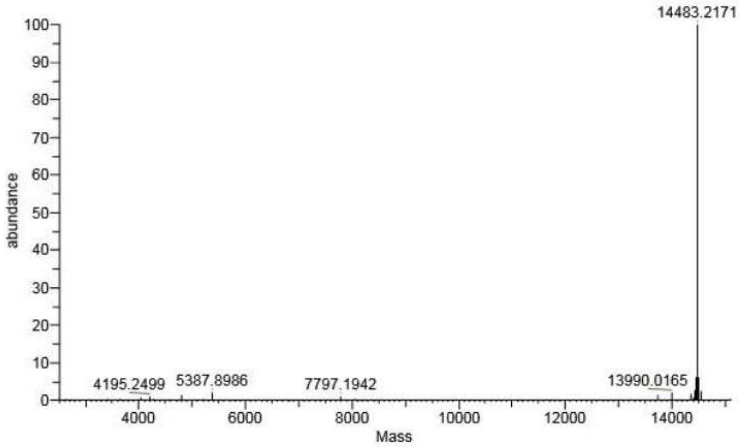

**Figure S8.** The ESI-MS spectrum of αSyn-140C. Theoretical mass 14492.13, experimental mass 14483.2171.

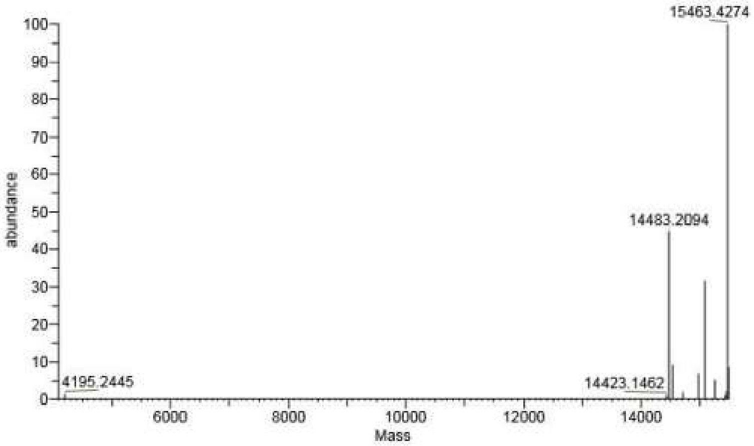

**Figure S9.** The ESI-MS spectrum of αSyn-AF647. Theoretical mass 15484.35, experimental mass 15463.42.

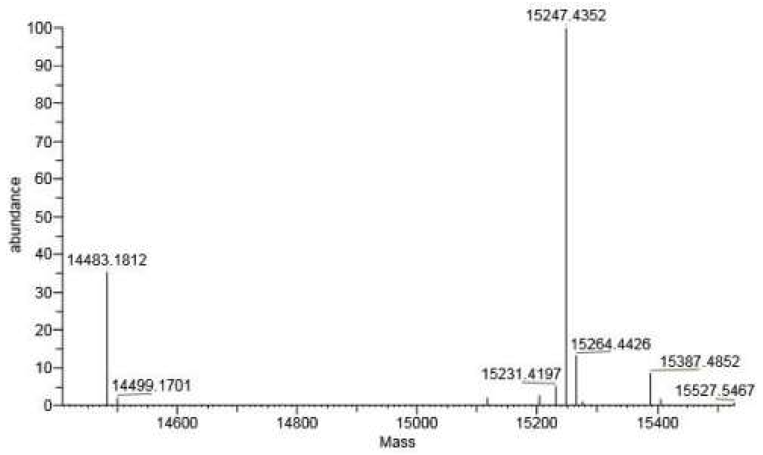

**Figure S10.** The ESI-MS spectrum of αSyn-dsCy5. Theoretical mass 15255.38, experimental mass 15247.44.

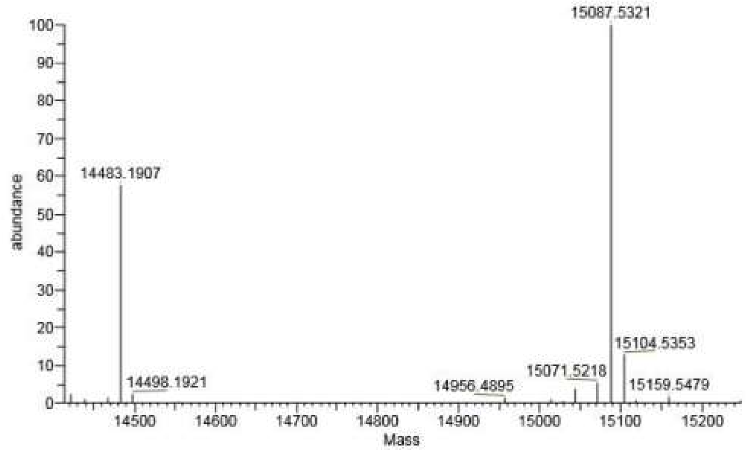

**Figure S11.** The ESI-MS spectrum of αSyn-Cy5. Theoretical mass 15097.48, experimental mass 15087.53.
